# A new experimental system to study meiotic recurrent non-allelic homologous recombination in budding yeast

**DOI:** 10.1101/2020.06.25.164855

**Authors:** Hailey N. C. Sedam, Juan Lucas Argueso

## Abstract

In humans, *de novo* recurrent copy number variations (CNVs) often arise during meiosis from non-allelic homologous recombination (NAHR) between low copy repeat elements (LCRs). These chromosomal rearrangements are responsible for a wide variety of genomic disorders involving duplication or deletion of dose-sensitive genes. The precise factors that steer meiotic cells toward this detrimental recombination pathway are not fully understood. To create a model for the investigation of LCR-mediated CNV mechanisms, we developed a diploid experimental system in *Saccharomyces cerevisiae*. We modified the right arm of chromosome V through the introduction of engineered LCRs: duplicated 5 to 35 kb segments of yeast DNA flanking single copy interstitial spacers, analogously to the meiotic NAHR substrates that exist in humans. Phenotypic markers, including a copy number reporter, were inserted within the interstitial spacer. Their segregation in the haploid meiotic progeny was used to phenotypically identity and classify recurrent CNV events. This system allowed us to measure the effects of LCR size on the frequency of meiotic *de novo* recurrent CNV formation, and to determine the relative proportions of each of the three main NAHR classes: interhomolog, intersister, and intrachromatid. The frequency of CNV increased as the LCRs became larger, and interhomolog NAHR was overrepresented relative to the two other classes. We showed that this experimental system directly mimics the features of *de novo* recurrent CNVs reported in human disease, thus it represents a promising tool for the discovery and characterization of conserved cellular factors and environmental exposures that can modulate meiotic NAHR.

## INTRODUCTION

Genomic disorders are diseases caused by structural rearrangements of human chromosomes (Lupski, 1998; Harel and Lupski, 2018). Such rearrangements are often the result of meiotic non-allelic homologous recombination (NAHR) leading to *de novo* recurrent copy number variation events (CNVs) in which a dose-sensitive disorder locus is duplicated or deleted (Carvalho and Lupski, 2016). Meiotic homologous recombination (HR) normally occurs between allelic sequences of properly aligned homologous chromosomes, sometimes producing reciprocal crossovers, but not structural rearrangements (Fig. 1A). Meiotic HR can also occasionally occur between misaligned non-allelic repetitive DNA substrates, such as the large low copy repeats (LCRs; >10 kb long, >97% identical nucleotide sequence) that are present in the human genome. In such cases, however, crossover resolution of HR intermediates leads to CNV formation (Liu et al., 2012; Kim et al., 2016). There are three primary ways through which meiotic NAHR between directly-oriented LCRs can generate recurrent CNVs. These are classified according to the position of the two recombination substrates relative to each other. Specifically, interhomolog NAHR (Fig. 1B) involves LCRs present one in each of the two homologous chromosomes; Intersister NAHR (Fig. 1C) involves LCRs present one in each of the two sister chromatids of the same homolog; and Intrachromatid NAHR (Fig. 1D) involves two LCRs present in the same chromatid.

**Figure 1.**
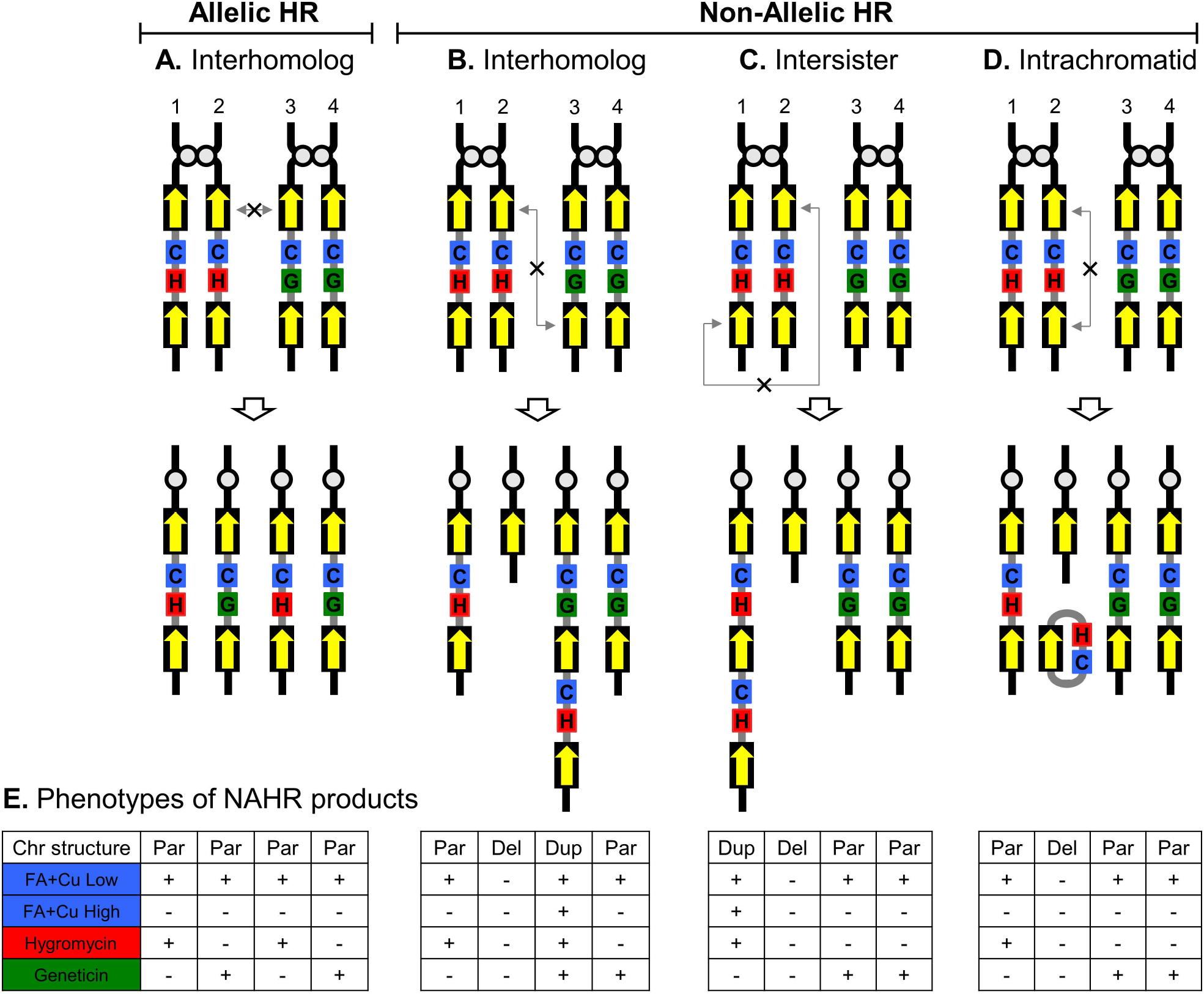
Meiotic recombination and phenotypic classification of recurrent NAHR events. (A-D) Schematic representation of the two replicated diploid Chr5 homologs at the start of meiotic recombination, not to scale. Four figures at the bottom of each panel represent the sets of four sister haploid cells after meiotic recombination. The black boxes with yellow arrows inside correspond to the LCRs. Thin gray lines connecting the LCRs and an X symbol indicate homologous recombination associated with a crossover outcome. The red H represents the *Hph*MX4 marker conferring resistance to Hygromycin B, the green G represents *Kan*MX4 marker conferring resistance to Geneticin. The blue C represents the dosage-dependent *SFA1^V208I^-CUP1* reporter cassette which, when duplicated, confers hyper-resistance to formaldehyde plus copper sulfate (FA+Cu). (E) shows the expected growth phenotype patterns of the spore colonies (from A-D) shown immediately above. +, indicates growth in media containing low concentration of FA+Cu, high concentration of FA+Cu, resistance to Hygromycin B, and resistance to Geneticin; -, indicates lack of growth.

Early studies established the presence of LCRs flanking critical genomic disorder loci as a necessity for recurrent *de novo* CNVs to form (Lupski, 1998, 2009; Carvalho and Lupski, 2016). Later on, the relative NAHR frequency was found to be modulated up or down by variations in locus architecture (*i.e*., LCR number, size, distance, orientation, and identity) (Liu et al., 2011b; Verges et al., 2017), as well as by the propensity for meiotic DSB formation within the LCRs mediated by PRDM9 ([MIM 609760]) (Borel et al., 2012; Pratto et al., 2014).

In 2011 the Lupski group explored the reciprocal recurrent CNVs that cause Smith-Magenis microdeletion syndrome (SMS [MIM 182290]) and Potocki-Lupski microduplication syndrome (PTLS [MIM 610883]) on chromosome 17p11.2 to specifically investigate the effect of LCR size and distance on NAHR frequency (Liu et al., 2011b). This region provides an excellent model for the analysis of recurrent CNV formation in humans because both the reciprocal duplication and deletion cause defined disease phenotypes, and a cohort of patients with well-characterized *de novo* CNV structures is available (Liu et al., 2011b). In addition, the dose-sensitive gene in this region (RAI1 [MIM 607642]) is flanked by at least three different pairs of LCRs that vary in size and interstitial distance. By determining the relative abundance in the patient cohort of recurrent *de novo* CNVs with boundaries at each of the three LCR pairs the authors were able to identify a strong positive correlation between LCR size and NARH frequency (R^2^=0.85627). In other words, more patients carried recurrent CNVs mediated by the longer LCRs, whereas fewer patients had CNVs with endpoints at the shorter LCRs.

Although it is clear that locus architecture and presence of recombination hotspots within LCRs are key determinants of recurrent NAHR frequency, other undiscovered factors may also exist. A single-sperm PCR-based genotyping study showed that recurrent NAHR frequency was variable at the *CMT1A-REP* locus within a cohort of men with the same LCR architecture (MacArthur et al., 2014). However, and strikingly, the frequency of NAHR was very similar within pairs of monozygotic twin brothers. Variation in genotypes at PRDM9 could not explain the variation in NAHR frequency, thus indicating that additional genetic and/or environmental factors likely contributed to recurrent CNV risk.

These unknown factors might include enzymes involved in HR or coordination of chromosome pairing (Goldman and Lichten, 1996; Grushcow et al., 1999; Goldman and Lichten, 2000; Shinohara and Shinohara, 2013; Shinohara et al., 2019), as well as exposure to chemicals that may possibly perturb some of these key meiotic processes (Shin et al., 2019; Cuenca et al., 2020). Therefore, sensitive cell-based assay systems able to recapitulate the LCR-mediated meiotic NAHR process experimentally are needed in order to identify the genes and environmental factors that can modulate recurrent CNV formation (Lupski, 2015; Yauk et al., 2015; Conover and Argueso, 2016). In this study, we describe the development of a one such assay in budding yeast cells. We show that this system allows direct and accurate phenotypic identification of meiotically-derived haploid cells carrying *de novo* recurrent duplications and deletions mediated by NAHR between engineered LCRs. We applied this assay to show experimentally that LCR size does indeed strongly correlate with CNV formation frequency, thus mimicking the behavior of the human 17p11.2 region. This result validated the use of this yeast model as a germane and promising approach to dissect the fundamental, and likely conserved, mechanisms that govern meiotic *de novo* recurrent CNV formation in eukaryotes.

## RESULTS and DISCUSSION

### Construction of the recurrent CNV reporter locus

Our meiotic CNV assay system consisted of a series of *Saccharomyces cerevisiae* strains harboring an engineered chromosomal locus with LCRs of varying sizes flanking phenotypic markers that allowed detection and classification of recurrent CNVs produced by meiotic NAHR (Fig. 2). We started with a *MATα* haploid strain (FCR8; Formaldehyde-Copper Resistant clone #8) containing a 59.6 kb tandem segmental duplication on the right arm of chromosome V (Chr5) that resulted from mitotic NAHR between two Ty1 retrotransposon element insertions (Fig. 2A-C) (Stanton, 2012). We then used a PCR-mediated targeted deletion approach (Goldstein and McCusker, 1999) to knock out the centromere-proximal (left side) portion of the proximal duplicated segment in FCR8, resulting in a Chr5 configuration with two identical 35 kb directly-oriented engineered LCRs separated by a 12 kb single copy interstitial spacer (IS) region containing the *SFA1^V208I^-CUP1-Kan*MX4 copy number reporter cassette (Fig. 2A-B) (Zhang et al., 2013; Klein et al., 2019). These 35 kb LCRs correspond to regions 2-5 in Fig. 2, and the single copy IS corresponds to region 1.

**Figure 2.**
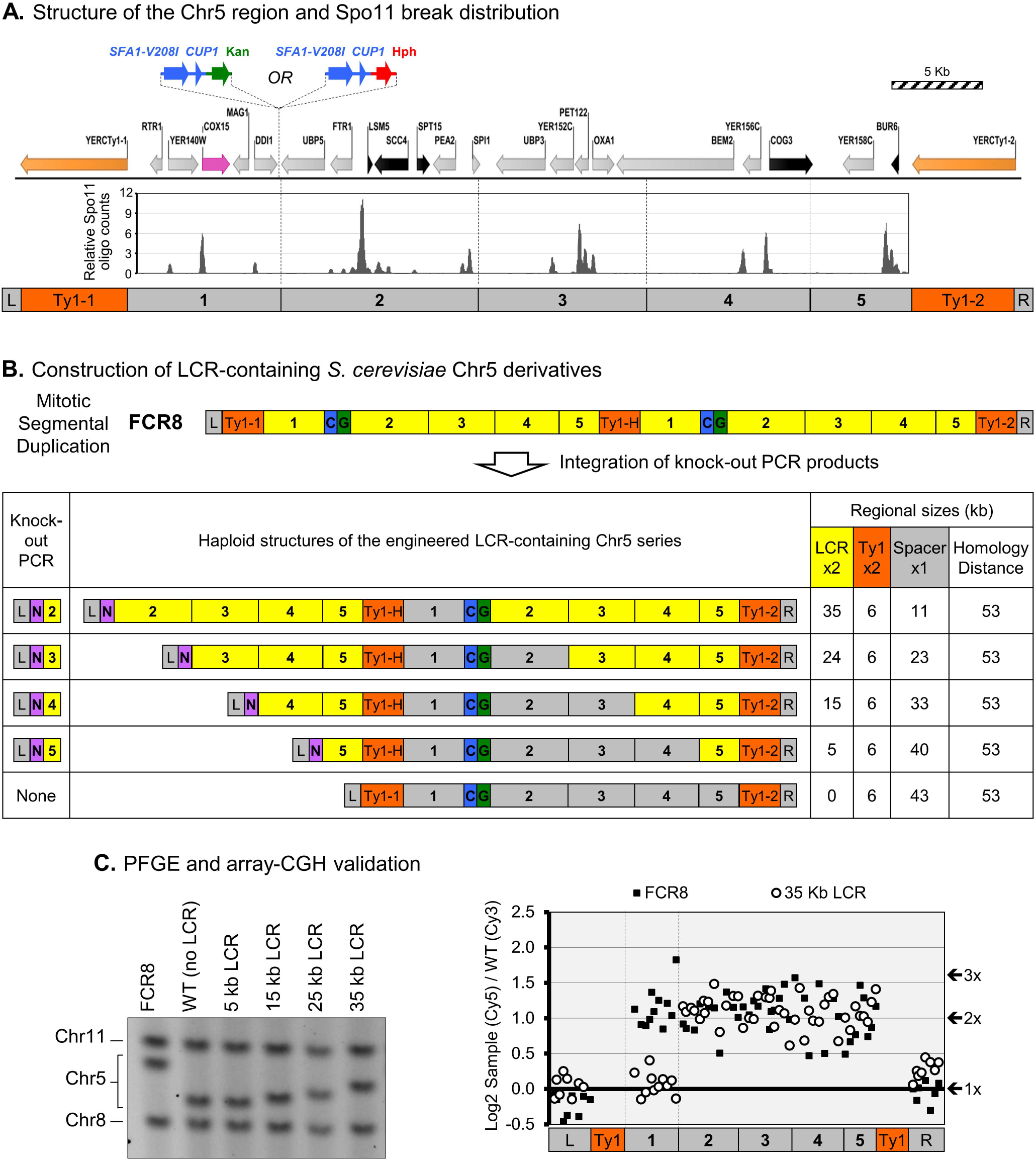
Construction of engineered LCRs in yeast chromosomes. (A) Structure of the *S. cerevisiae* Chr5 region analyzed and Spo11 DSB distribution within it. Orange arrows indicate the retrotransposon elements *YERCTy1*- (left) and *YERCTy1*-2 (right). Blue arrows indicate the *SFA1^V208I^-CUP1* dosage dependent reporter cassette. Green and red arrows indicate Geneticin and Hygromycin B resistance genes, respectively. Black arrows indicate genes essential for viability (their deletion is lethal). The pink arrow indicates the *COX15* gene (its deletion is viable, but causes a *petite* phenotype). Gray arrows indicate all other genes present in this experimental region. Peaks in the plot below indicate the specific positions where Spo11 creates meiotic double strand breaks that initiate meiotic recombination. The assay region was divided in regions 1-5, according to the boundaries used in the LCR construction. (B) Construction of LCR-containing Chr5 derivatives. The structure of Chr5 in FCR8 is shown. L and R indicate the left (centromeric) and right (telomeric) boundaries of the duplicated region. Orange boxes indicate Ty1 elements. Ty1-H indicates the hybrid *YERCTy1*-1 / *YERCTy1*-2 rearrangement junction structure present in FCR8. Yellow boxes indicate duplicated segments, gray boxes indicate the interstitial spacer area, blue boxes indicate the *SFA1^V208I^-CUP1* dosage dependent reporter cassette, green boxes indicate resistance to Geneticin (the matching homolog constructs were resistant to Hygromycin B instead). The purple N the indicates Nourseothricin (*Nat*MX4) resistance marker that was used to knock out the proximal portion of the FCR8 duplicated region to produce LCRs. LCR, Ty1, spacer, and homology distance sizes are provided in kb. (C) PFGE and array-CGH validation of LCR-containing Chr5 derivatives. Chr11 (665 kb) and Chr 8 (540 kb) are also shown as size references; the upper and lower sections of the PFGE were cropped out to emphasize the region of interest. The array-CGH plot to the right represents the copy number of array probes in the experimental region of Chr5. Individual data points correspond to the Log2 of the (Cy3-labeled sample DNA / (Cy5-labeled reference DNA signal (Y-axis) indicative of relative copy number for each probe, plotted over the Chr5 coordinate of each probe (X-axis). The reference DNA in this experiment was from the wild type haploid strain from (C), without any LCRs (single copy for regions L, 1-5, R). The predicted discrete copy number levels (1x, 2x, 3x) are indicated to the right. The positions of the relevant Chr5 segments from (A) are indicated in the X-axis. The symbol legends for the FCR8 strain and the parent 35 kb LCR strain are shown above the plot. No other copy number deviations from 1x were detected anywhere else in the genome.

We validated the structure of the 35 kb LCR strain using pulse field gel electrophoresis (PFGE) and microarray-based comparative genomic hybridization (array-CGH) (Fig. 2C). PFGE revealed a downward shift of Chr5 in the 35 kb LCR strain when compared to FCR8, corresponding to loss of one copy of region 1. Array-CGH further confirmed that regions 1-5 were duplicated in FCR8, but in the 35 kb LCR strain region 1 had reverted back to single copy while regions 2-5 remained duplicated.

We next switched the mating type of the haploid containing the 35 kb LCR Chr5 to *MATα*, and swapped the *Kan*MX4 marker for *Hph*MX4. The *Kan*MX4 and *Hph*MX4 markers confer resistance Geneticin (Gen) or Hygromycin B (Hyg), respectively (Wach et al., 1994; Goldstein and McCusker, 1999). The *SFA1^V208I^-CUP1* construct is a gene dosage sensitive reporter that when duplicated enable cells to grow on media containing high concentrations of formaldehyde plus copper sulfate (FA+Cu). Cells carrying the parental configuration of Chr5, with a single copy of the reporter are resistant to low FA+Cu concentrations, while cells with a deletion of this reporter are sensitive. These different growth patterns can be used to determine the copy number of the reporter cassette, and thus detect CNVs. Finally, the *MATα* and *MATα* haploids were mated to each other to create an experimental diploid which was homozygous for the flanking 35 kb LCRs and the *SFA1^V208I^-CUP1* copy number reporter, but hemizygous for either *Kan*MX4 or *Hph*MX4 (Fig. 1A-D).

### Experimental detection and characterization of recurrent CNVs

We induced meiosis of the diploid 35 kb LCR diploid strain, leading to the formation of the characteristic yeast tetrad asci that contain the four haploid spore progeny of a single meiotic division. These haploid spores are analogous to human sperm or eggs, but they remain grouped together inside the ascus, which can be micro-dissected to separate and culture each spore cell individually. This allows the recovery of a defined group of four colonies derived from the sibling cells, and their subsequent genotypic and phenotypic characterization can then be interpreted in the context of a single complete meiotic cell division. The diploid strain used in our assay was designed so that allelic recombination and each of the three NAHR classes conferred a different combination of FA+Cu, Gen and Hyg resistance phenotypes between sibling cells within each tetrad (Fig. 1E).

We micro-dissected 323 tetrads derived from the 35 kb LCR diploid in rich medium (YPD) so that each haploid spore was allowed to germinate and grow mitotically into a colony. These were then replica-plated to phenotypically assess the presence of CNVs. Overall spore viability was high (90.2%), yielding 245 tetrads in which the complete set of four sibling spores germinated and formed a colony. Lack of recombination or normal allelic recombination produced cells with either parental phenotypes that displayed resistance to one of the two drugs (never both together), and low FA+Cu resistance. For the purpose of this assay, we diagnosed CNVs as the loss or gain of the reporter cassette where a loss (deletion) leads to sensitivity to Gen, Hyg and FA+Cu, and a gain (duplication, or rarely triplication) leads to the ability to grow on media containing a high concentration of FA+Cu, and a specific pattern of drug resistance depending on the NAHR class (Fig. 1E). Interhomolog NAHR produced tetrads with a pair of parental phenotype cells displaying either drug resistance and resistance to low FA+Cu concentrations, one cell carrying a duplication displaying resistance to the two drugs and to a high concentration of FA+Cu, and one cell carrying the reciprocal deletion with double drug sensitivity and sensitivity to low FA+Cu (Fig. 1B, E). As an additional phenotype associated with deletions specifically in the 35 kb LCR constructs, loss of the *COX15* gene present in the IS region (pink arrow, Fig. 2A) rendered colonies smaller than normal (*petite* phenotype) in YPD, and unable to grow on media containing a non-fermentable carbon source (YPGE). Intersister NAHR produced tetrads containing a pair of parental phenotype cells resistant to the same drug, one cell containing the duplication with resistance to high FA+Cu and the drug resistance opposite of the two parental sibling cells, and one cell containing the reciprocal deletion with sensitivity to both drugs, low FA+Cu, and *petite*. Finally, intrachromatid NAHR produced tetrads containing a pair of parental phenotype cells with single drug resistance of the same type, another parental phenotype cell with single drug resistance of the opposite type, and one cell containing a deletion leading to sensitivity to both drugs, low FA+Cu, and *petite* (Fig. 1D). The reciprocal product of intrachromatid NAHR in this case was a circular DNA molecule that does not contain any origins of DNA replication nor a centromere, thus it was incapable of propagation during the growth of the colonies and did not contribute to phenotype.

To validate our phenotypic analyses, we randomly selected tetrads from which NAHR had been phenotypically classified as interhomolog, intersister, or intrachromatid, and characterized colonies from all four sibling spores via PFGE, array-CGH, and quantitative digital droplet PCR (ddPCR) (Fig. 3). The parental Chr5 with 35 kb LCRs migrates in PFGE to a position between Chr8 and Chr11 (Fig. 3A). As expected, the tetrads derived from the interhomolog and intersister NAHR classes had two colonies with parental sized Chr5, one contained a longer Chr5 due to a duplication, and one contained a shorter Chr5 due to a deletion. The intrachromatid class showed three colonies that contained parental sized Chr5, and one contained a deletion within Chr5, without a reciprocal duplication. We analyzed one parental, one duplication, and one deletion containing haploid from one interhomolog tetrad using array-CGH and confirmed the presence of the expected Chr5 copy-number probe signal predicted by phenotype and PFGE (Fig 3B). The parental spore colony had signal equivalent to a single copy of the IS region 1 and two copies of the LCRs in region 2-5, the duplication colony had two copies of the IS region 1 and three copies of the LCR regions 2-5, and the deletion colony had lost all signal from the region 1 probes (zero copies of the IS region 1) and only one copy of the LCR region 2-5. Finally, we also performed ddPCR using primers for the *SFA1, Kan*MX4, *Hph*MX4 markers-all of which were inside the IS region 1, and for a centromere-proximal control region of Chr5 outside the engineered segment and thus predicted to not be involved in the NAHR events (Fig. 3C; Methods). We found that in the parent strain and in all parental spores the proximal control region, *SFA1*, and either *Kan*MX4 or *Hph*MX4 were present at one copy as expected. All deletions had a single copy of the proximal region, and complete loss of signal from the IS ddPCR markers. The interhomolog duplication had two copies of *SFA1*, and one copy of each *Kan*MX4 and *Hph*MX4. In contrast, the intersister duplication had two copies of *SFA1* and two copies of *Kan*MX4, but zero copies of *Hph*MX4. Taken together, these analyses confirmed that for each meiotic NAHR class the growth phenotypes of the spores directly reflected their respective Chr5 structural configurations (copy number gain or loss of the IS and LCR regions). No additional chromosomal rearrangements were detected elsewhere in the genomes of these spore colonies by PFGE or array-CGH, indicating that CNVs were confined to the experimental chromosomal region engineered for the NAHR assay.

**Figure 3.**
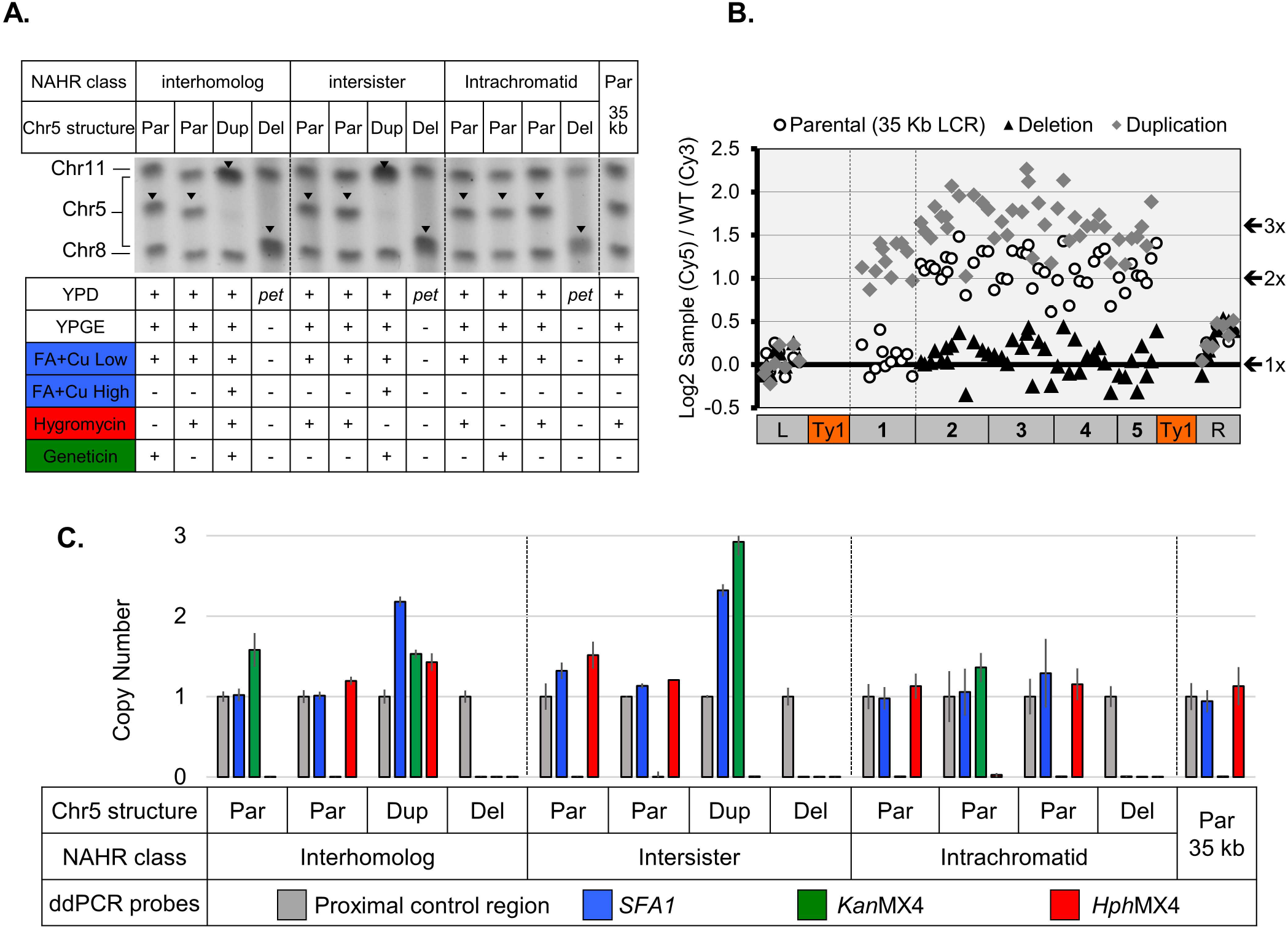
Phenotypic and molecular characterization of meiotic NAHR in the 35 kb LCR strain. All items represented in the same way as in Figs. 1–2, with noted exceptions. (A) PFGE of the four haploid spores from one tetrad from each class of NAHR. The phenotypes of each spore colony are indicated below the PFGE lane. +, growth; -, no growth; *pet*, *petite*. Par, parental phenotype and Chr5 size; Dup, duplication of the CNV region; Del, deletion of the CNV region. Black inverted triangles indicate the Chr5 band. (B) array-CGH of a parental phenotype spore, a duplication containing spore, and a deletion containing spore. (C) Plot of ddPCR results measuring normalized copy number of *SFA1^V208I^, Hph*MX4, and *Kan*MX4 as well as of a proximal region not involved in the rearrangements used for normalization. The data are presented for DNA prepared from the corresponding samples represented in PFGE (A).

It is well established that there is an intrinsic bias favoring the use of the homolog as a donor template for repair of double strand breaks during allelic meiotic recombination. This is believed to be necessary for establishing physical connections between homologs (*i.e*., chiasmata) which are essential for proper chromosomal disjunction of during the first meiotic division (Schwacha and Kleckner, 1997). However, it is still unclear whether this bias also applies during non-allelic meiotic recombination, for example between LCRs. This is an extraordinarily difficult problem to study in *de novo* recurrent CNV human genomic disorders, as it requires phased haplotype information from the chromosomes of the healthy parents and of the affected patient. In one example, when haplotype data at the 16p11.2 locus was restricted to high confidence haplotypes, there appeared to be a trend towards the presence of an interhomolog NAHR bias (Duyzend et al., 2016). However, when all phasing data was combined, the trend toward interhomolog NAHR was abolished. We investigated the presence of a bias within our system by comparing the relative observed abundance of tetrads containing CNVs from each of the three NAHR classes to the frequency expected if the selection of a repair template (donor) sequence were random. If no bias were present, the interhomolog NAHR proportion should be 50% (there are two non-allelic recombination donors are available in the homolog), whereas intersister and intrachromatid NAHR proportions should be 25% each (one non-allelic donor available in the sister chromatid, and one non-allelic donor in the same chromatid). In the 35 kb LCR experimental diploid strain we measured the NAHR proportions at 49 (63.6%) interhomolog : 17 (22.1%) intersister : 11 (13%) intrachromatid. This observed distribution was statistically different from the neutral 2:1:1 ratio expected if non-allelic LCR donor choice were random (X^2^= 6.6623, p-value = 0.03575). This result suggests that, at least in this system and for the 35 kb LCR configuration, NAHR interactions are biased toward interhomolog at the expense of intrachromatid events. A similar trend was later observed for shorter LCR constructs (see below; Fig. 5C).

While the three NAHR classes shown in Fig. 1 are the most abundant, there is also evidence for rare *de novo* recurrent tandem triplication (and other even more complex CNVs) of dose-sensitive genes in human genomic disorders (Liu et al., 2011a; Harel and Lupski, 2018). Interestingly, we were able detect phenotypically five examples of double-NAHR leading to triplication events within single meiotic divisions. We characterized one of these tetrads by PFGE and array-CGH (Fig. 4) to confirm the structural rearrangements. PFGE of the four sibling spores from a single tetrad showed that one contained a parental sized Chr5, one contained a much longer Chr5 consistent with a triplication, and two contained a deletion on Chr5 (Fig. 4A). We further confirmed the triplication through array-CGH which showed that this spore contained three copies of the IS region 1, and four copies of the LCR regions 2-5 (Fig. 4B-C). Additionally, we identified three tetrads with complex patterns of marker segregation consistent with pre-meiotic mitotic chromosomal rearrangements (data not shown). Even though these presumably mitotic events were detected, their frequency was relatively low (<1%), therefore they were not likely to have interfered significantly with the accuracy of meiotic NAHR frequency measurements.

**Figure 4.**
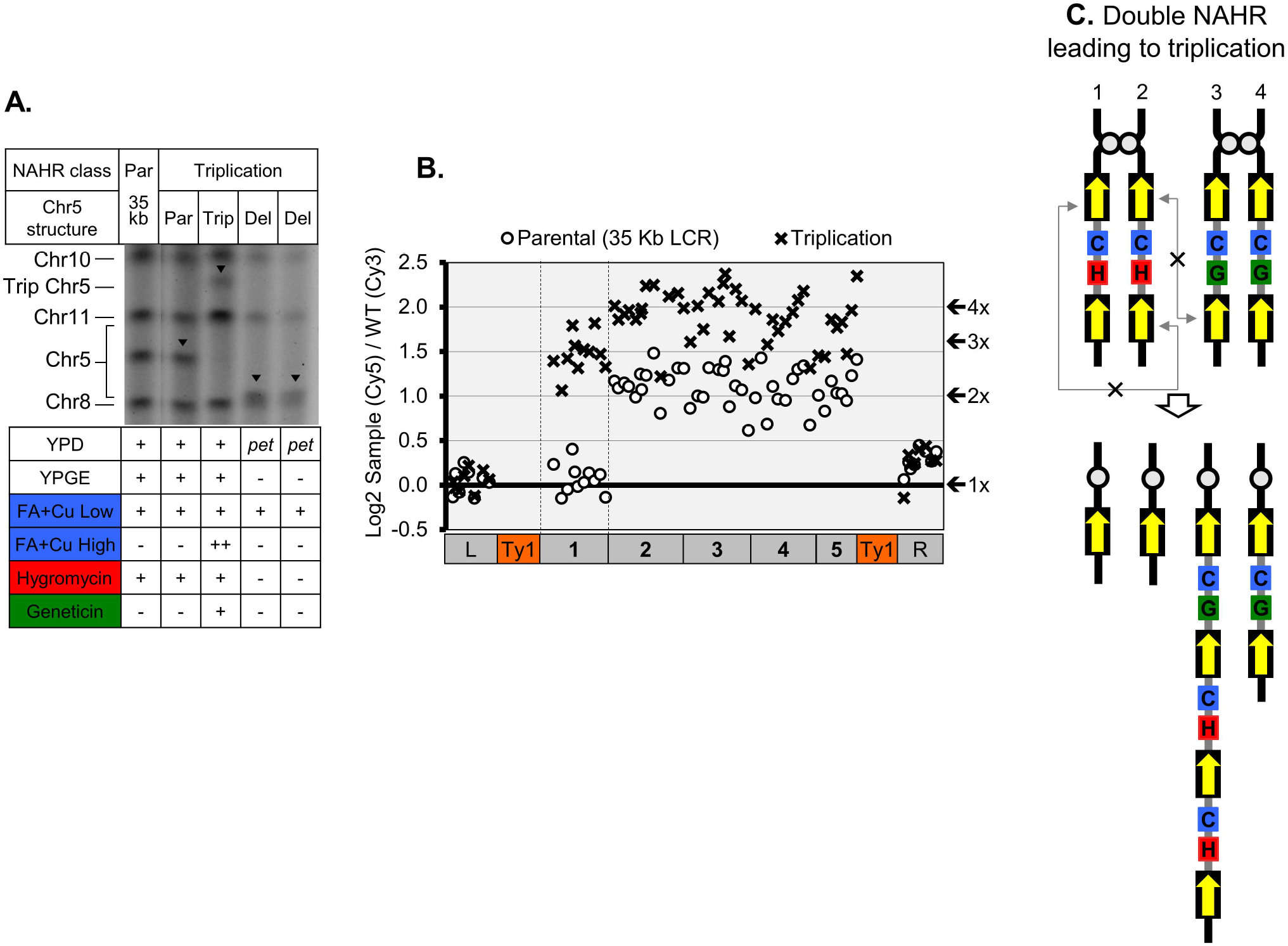
Characterization of a triplication CNV event All items represented in the same way as in Figs. 1–3, with noted exceptions. (A) PFGE and phenotypes of four spores recovered from a single tetrad of the 35 kb LCR strain containing one parental, one triplication, and two deletion NAHR events. Trip, Chr5 containing the triplication event. (B) array-CGH plot shows a parental phenotype spore and the spore containing a triplication event from (A). (C) Schematic representation of one of the ways through which double NAHR can lead to the triplication outcome detected in this example.

The combined results from the 35 kb LCR strain demonstrated that our experimental model system allows for faithful phenotypic identification of recurrent CNVs of all three meiotic NAHR classes. In addition, this initial analysis suggested that meiotic NAHR is likely subject to the same interhomolog bias that is well described for meiotic allelic HR, and also showed that this approach is capable of detecting complex and rare recombination events such as triplications and pre-meiotic CNVs.

### Relationship between LCR size and CNV frequency

As an initial application of this experimental approach, we decided to revisit the question of whether the size of LCRs can influence CNV frequency, and if so, if a correlation could be traced analogously to that shown to exist at the human SMS and PTLS 17p11.2 locus (Liu et al., 2011b). To do so, we used a similar chromosome engineering approach to the one used to create the 35 kb LCR strain. In this case, we progressively increased the size of the PCR-targeted deletion of the duplicated segment from FCR8, which allowed for the creation of varied LCR sizes, while maintaining a constant ~53 kb distance between homologies (Fig. 2B). The meiotic recombination hotspots in *S. cerevisiae* have been thoroughly mapped via sequencing of terminal ssDNA fragments that remain attached to the conserved meiotic recombination initiator Spo11 (Pan et al., 2011). Using this map as a general guide, we designed shorter LCRs such that they would all contain at least one predicted meiotic DSB hotspot (Fig. 2A). Though each LCR also contained a ~6 kb Ty1 element at their distal end, it is important to note that the recombination properties of yeast Ty1 elements have been studied previously and shown to be repressed for meiotic recombination (Kupiec and Petes, 1988). Accordingly, we saw very low CNV occurrence (<0.2%) in a control strain with the normal Chr5 configuration (contained only the flanking Ty1s, but lacked engineered LCRs). Since the Ty1 insertions behaved essentially as inert sequences, each LCR-containing strain was identified by the length of their respective unique DNA (regions 2-5), not including the length of the Ty1 elements. We produced four additional experimental strains through this approach: 0 kb (control strain; Ty1 only), 5 kb, 15 kb, and 24 kb LCRs (Fig. 2 B-C). We then switched the mating type and drug resistance marker of each of the LCR containing haploids and mated them to create diploids which were homozygous for each flanking LCR size and the *SFA1^V208I^-CUP1* copy number reporter, but hemizygous for either *Kan*MX4 or *Hph*MX4.

Approximately 300 tetrads were dissected for each new LCR diploid strain and the resulting spore colonies were phenotypically scored for the presence of CNV duplications (Fig. 5). The large proximal deletions required to create the smaller LCRs had the consequence of moving genes essential for yeast viability into the IS region (black arrows in regions 2, 4, 5 in Fig. 2A). Therefore, unlike the 35 kb LCR described earlier, absence of the IS region in spores carrying the deletion CNVs led to inviability (Fig. 2A-B). Due to the inability to grow colonies carrying lethal deletions, in these new LCR strains we were unable to positively call CNVs using whole tetrad (4-spore) information. However, we were still able to recover viable spores carrying duplications, and detected them phenotypically through resistance to high concentrations of FA+Cu. In addition, we were still able to sub-categorize these, based on the presence of double drug resistance (Gen and Hyg) for interhomolog duplications, versus single drug resistance (Gen or Hyg) for intersister duplications (Fig. 1B-C, E). We performed PFGE karyotypic validation in 15 kb LCR strain by randomly selecting two 3-viable spore tetrads, one with presumed interhomolog and one with presumed intersister duplication events (Fig. 5A). We observed two spores with parental Chr5, and the expected longer Chr5 band in both the interhomolog and intersister duplication containing colonies, indicative of duplication CNVs (Fig. 5A). Additionally, ddPCR revealed a single copy of the proximal control, *SFA1* and *Kan*MX4 or *Hph*MX4 markers in all parental phenotype colonies and a duplication of *SFA1* in both duplication colonies, with a single copy of each *Kan*MX4 and *Hph*MX4 in the interhomolog duplication, and two copies of *Hph*MX4, but absence of *Kan*MX4, in the intersister duplication (Fig. 5B). Therefore, these molecular analyses again confirmed that the FA+Cu and drug resistance phenotypes were directly indicative of the structural configuration present in the genome.

**Figure 5.**
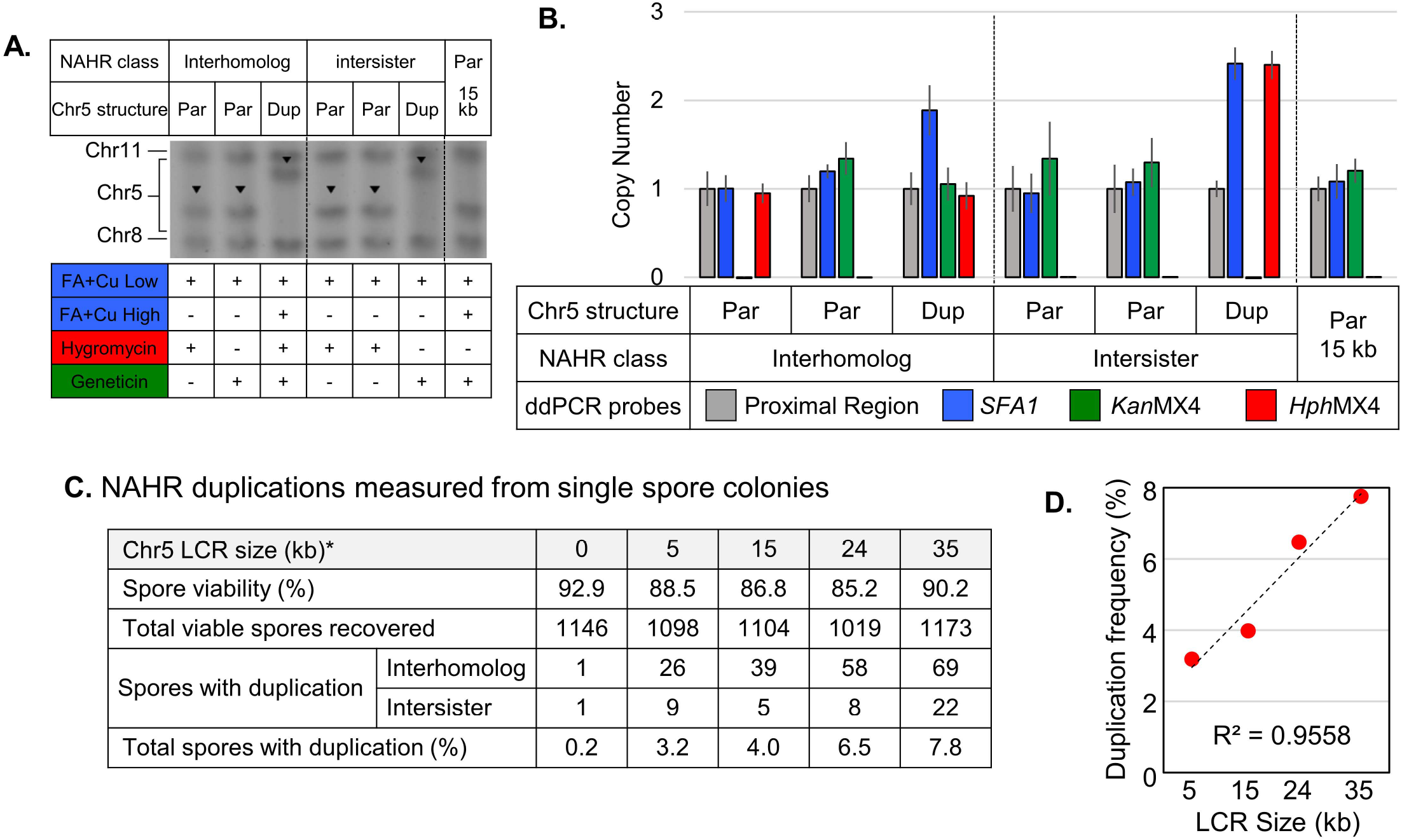
Structural validation of duplications derived from the 15 kb LCR strain, and CNV duplication analyses. All items represented in the same way as in Figs. 1–4, with noted exceptions. (A) PFGE karyotypes showing representatives of interchromosome and interchromatid NAHR from the 15 kb LCR strain. Phenotypes for each spore are indicated below the PFGE lanes. (B) Copy number plots from ddPCR results. Data correspond to the same spores represented in (A). (C) Spore viability and NAHR duplication frequency data from the engineered LCR strain series. (D) Linear regression of data from (C) where the X-axis is LCR size and the Y axis is (interhomolog + intersister) duplication CNV frequency.

Next we compared the total spore viability, frequency, and classes of duplications detected in the LCR strain set. We observed a slight, yet distinctive trend in spore viability among the four experimental diploids (0, 5, 15, and 24 kb LCR) where deletions were lethal. Spore viability was highest for the diploids with no LCR, and progressively decreased as the LCRs got longer. This suggested that meiosis in diploids with longer LCRs produced more inviable spores, presumably those carrying lethal deletions. Notably, spore viability was rescued (90.2%) in the longest LCR construct (35 kb), likely because deletions were viable in that strain (Fig. 3).

Finally, we counted the spore colonies carrying duplications and interrogated the presence of a biased HR template repair choice, as well as their overall frequencies in the engineered LCR strain series. Interhomolog duplications were more than twice as abundant as intersister duplications in all LCR strains, and when combined (192:44, respectively) were significantly overrepresented relative to the neutral 2:1 ratio expected for random donor choice (X^2^=22.938; p=0.0001). This result was consistent with the presence of interhomolog bias detected earlier through whole-tetrad data in the 35 kb strain. We also interrogated the data to determine whether a positive correlation existed between LCR size and duplication frequency. We calculated the total (interhomolog plus intersister) duplication frequency for each LCR size and detected a clear association. Analogously to the Liu *et al*. 17p11.2 findings, the data showed a significant positive linear correlation between the size of the LCRs and the frequency of recurrent duplication CNVs (Fig. 5D; R^2^=0.9558, p-value=0.02235). Therefore, our experimental results in model yeast cells directly agreed with the clinical observation of larger LCRs promoting higher frequencies of recurrent CNV formation.

### Conclusions and future perspectives

The results from this study showed that meiotic NAHR operates similarly in budding yeast and in humans, thus providing an experimental model for further studies. In recent years the need for such an assay system has become urgent, as evidence suggests that the factors involved in meiotic recurrent and mitotic non-recurrent CNV formation are likely different (Lupski, 2015; Conover and Argueso, 2016). Therefore, we are continuing to improve this assay system to allow for further probing of environmental and genetic factors that may influence NAHR. This will eventually provide a high throughput tool for the systematic investigation of the structural factors involved in meiotic NAHR such as LCR size, distance, identity, and relative position in chromosomes (*e.g*., near telomeres), as well as screening of candidate environmental toxicants and analysis of conserved enzyme activities that may be involved in the formation of meiotic *de novo* recurrent CNVs and their subsequent transmission to the offspring.

## MATERIALS and METHODS

### Yeast strains and plasmids

All *Saccharomyces cerevisiae* strains used in this study were derived from CG379 strain background containing noted locus-specific changes introduced by PCR-based transformation (Fig. 2, Table S1) (Morrison et al., 1991). The precursor strain for the engineered LCRs was the FCR8 clone, a copper and formaldehyde hyper-resistant strain that had a mitotically-derived tandem segmental duplication on the right arm of chromosome 5 (Chr5) mediated by recombination between *YERCTy1*-1 and *YERCTy1*-2. FCR8 was obtained by selection for spontaneous FA+Cu resistance from culture of a haploid carrying a single copy of the *SFA1^V208I^-CUP1-Kan*MX4 reporter cassette inserted downstream of *DDI1* on Chr5 (Fig. 2A-B) (Stanton, 2012).

Through PCR-based, homologous recombination-mediated deletion of the FCR8 proximal *SFA1^V208I^-CUP1* cassette with the *Nat*MX4 cassette (Goldstein and McCusker, 1999), we were able to generate strains with one copy of the *SFA1^V208I^-CUP1-Kan*MX4 cassette flanked by repeated regions of yeast DNA of varying sizes (Fig. 2A-B). The homology segments used to target the knockout PCR products are represented in Fig. 2B (left column) and their specific nucleotide sequences are shown in Table S2. By increasing the region knocked out by PCR, we were able to alter the size of the LCRs inversely with the size of the interstitial spacer (IS) while keeping the distance between homologous regions constant. We used this method to create four haploid LCR containing strains: 5 kb LCRs flanking a 41 kb IS with 52.9 kb separating LCR homologies; 15 kb LCRs flanking a 33 kb IS with 52.9 kb separating LCR homologies; 24 kb LCRs flanking a 23 kb IS with 52.9 kb separating LCR homologies; and 35 kb LCRs flanking a 12 kb IS with 52.9kb separating LCR homologies (Fig. 2B). We then switched the mating type of the LCR containing haploids. Next, we used a PCR based transformation to swap the *Kan*MX4 Geneticin (Gen) resistance marker for *Hph*MX4 Hygromycin B (Hyg) resistance marker in one homolog. We then mated the opposite resistance marker containing haploid strains so their resulting diploids were homozygous for the LCRs and the *SFA1^V208I^-CUP1* reporter and hemizygous for either *Kan*MX4 or *Hph*MX4.

### Culture media and CNV selection conditions

Yeast diploid cells were induced to sporulate by first growing overnight 5 mL liquid pre-sporulation media cultures (8 g/L yeast extract, 3 g/L peptone, 100 g/L glucose, 10 g/L complete drop out mix, and 5 g/L methionine). Next they were centrifuged, cell pellets were washed twice with sterile distilled water and half the culture was put into 4 mL of liquid sporulation media (1 g/L yeast extract, 10 g/L potassium acetate, 0.5 g/L glucose, 2.5 g/L complete drop out mix, and 3.8 g/L methionine). Cells were incubated in liquid sporulation media at 25C while shaking for four days. On the fourth day culture were removed from the shaker and kept at 25C for up to one week for tetrads to be dissected. Tetrads were dissected on YPD agar (rich medium) and incubated for 2 days at 30C. Plates were then replica plated onto YPD plus 400 mg/L Gen, 400 mg/L Hyg, YPGE, and SC supplemented with a complete drop-out mix and a range of concentrations of copper sulfate and formaldehyde (200 μM CuSO_4_ / 2.3 mM FA, 250 μM CuSO_4_ / 2.5 mM FA, 300 μM CuSO_4_ / 2.7 mM FA). FA+Cu concentrations were optimized for tetrad replica plating based on parameters described earlier (Zhang et al., 2013). Formaldehyde-containing plates were poured fresh the day before they were used. 1 M dilutions [101.5 μL of 37% by weight Formaldehyde methanol stabilized stock (Fisher) in 1148.5 μL sterile water] of formaldehyde were made and placed opposite copper sulfate inside an Erlenmeyer flask immediately before media was added to the flask and plates were poured. Cells were incubated on YPD + Gen and YPD + Hyg plates at 30 C for 2 days, FA+Cu plates were incubated at 30 C for 4 days.

### Analysis of NAHR-mediated CNVs

To detect *de novo* meiotic recurrent CNVs caused by chromosome architecture we needed reporters capable of differentiating the expected single copy of our region of interest versus a copy number loss or gain. Our group previously optimized a CNV assay allowing for detection of single copy increase of the *SFA1^V208I^-CUP1* reporter cassette in mitotic cells, where a single duplication of the *SFA1^V208I^-CUP1* reporter cassette allows for growth on media containing copper sulfate and formaldehyde (Narayanan et al., 2006; Zhang et al., 2013; Klein et al., 2019). Additionally, because NAHR can occur between LCRs on the homologous chromosome, the sister chromatid, or within the same chromatid (Fig. 1B-D), detection of the maternal allele versus the paternal allele was also needed for the classify these recurrent CNVs. By adding the either the *Kan*MX4 gene or the *Hph*MX4 gene to the *SFA1^V208I^-CUP1* reporter, we were able to differentiate between which LCRs the CNV causing NAHR occurred based on growth on Gen or Hyg containing media (Fig. 1E).

### Molecular karyotype analyses

Candidate sibling spores from each CNV phenotypic class were randomly selected for PFGE, digital droplet PCR and array-CGH to validate structural rearrangements present in them. Full length chromosomal DNA was prepared in agarose and was separated based on length via PFGE to reveal size changes in Chr5. Genomic DNA was prepared from PFGE agarose plugs as previously described and used for array-CGH using the methods described previously (Zhang et al., 2013). These DNA samples were also used for quantitative digital droplet (ddPCR). Sheared DNA was further fragmented by digestion with *Mfe*I-HF (New England Biolabs), a restriction enzyme that cuts frequently and between the engineered LCRs, but did not cut between ddPCR primers. Digested DNA was diluted to 0.05 ng/μL and 2 μL of this dilution was used as a template for each ddPCR reaction. The diluted DNAs were analyzed using a combination of four ddPCR primer sets, each for a specific region of Chr5. Specific ddPCR primer sequences are shown in Table S2. Biorad’s QX200 EvaGreen Supermix and protocol were used for all ddPCR reactions. For each experimental ddPCR reaction, the same DNA underwent ddPCR for a control single copy region directly proximal to the LCR. The signal from this proximal region was used to normalize the concentration of the DNA internally for each DNA template, allowing comparison of gene copy number with each experimental probe. One haploid parent was selected as a control for each ddPCR reaction and a no-DNA control was run with each experimental ddPCR reaction. Before normalization or standardization any residual fluorescence seen in the control was subtracted from all experimental values.

### Statistics employed

We performed a linear regression analysis on our CNV frequency versus LCR size using the lm function in R studio version 3.4.0. CNV frequency was calculated as the total number of spores containing *SFA1^V208I^-CUP1* duplications divided by the total number of living spores analyzed for each strain. Spore data did not include intrachromatid events as deletions were only viable in the 35 kb LCR strain. Additionally, we performed a Chi-Squared Test for Given Proportions on the ratios of each NAHR modality used within the 35 kb LCR strain using the chisq.test function in R studio version 3.4.0. Proportions were calculated as the number of complete tetrads containing an interhomolog, intersister, or intrachromatid recombination event versus the total number of complete tetrads containing recombination events.

## ACKNOWLEDGMENTS

We thank Jim Lupski, Claudia Carvalho, Tom Glover, Tom Petes, and Michael Lichten for valuable discussions and insights. JLA received research grants from Boettcher Foundation and from the Escher Fund for Autism during early phases of this project. Research reported here was subsequently supported by an NIH grant to JLA (R35GM119788).

---

**Supplemental Table S1.** Strains used in this study.

| LCR Size | *MAT* | *KanMX4*<br>haploids | *HphMX4*<br>haploids | Experimental<br>diploids |
| --- | --- | --- | --- | --- |
| FCR8 | $\alpha$ | JAY871 | - | - |
| 0 kb | a |  | JAY2008 | JAY2025 |
| | $\alpha$ | JAY2026 | | |
| 5 kb | a | JAY1952 |  | JAY2117, JAY2118 |
| | $\alpha$ | | JAY2113, JAY2114 | |
| 15 kb | a | JAY880, JAY881 |  | JAY1849, JAY1852 |
| | $\alpha$ | | JAY879 | |
| 24 kb | a | JAY1954 |  | JAY1965 |
| | $\alpha$ | | JAY1950 | |
| 35 kb | a |  | JAY1843 | JAY1910, JAY1911 |
| | $\alpha$ | JAY1888 | | |

The *MATa* and *MAT $\alpha$* haploids with the same LCR construct and either *KanMX4* or *HphMX4* markers were mated to form the corresponding experimental diploid strains used in the meiotic analysis.

**Supplemental Table S2.**
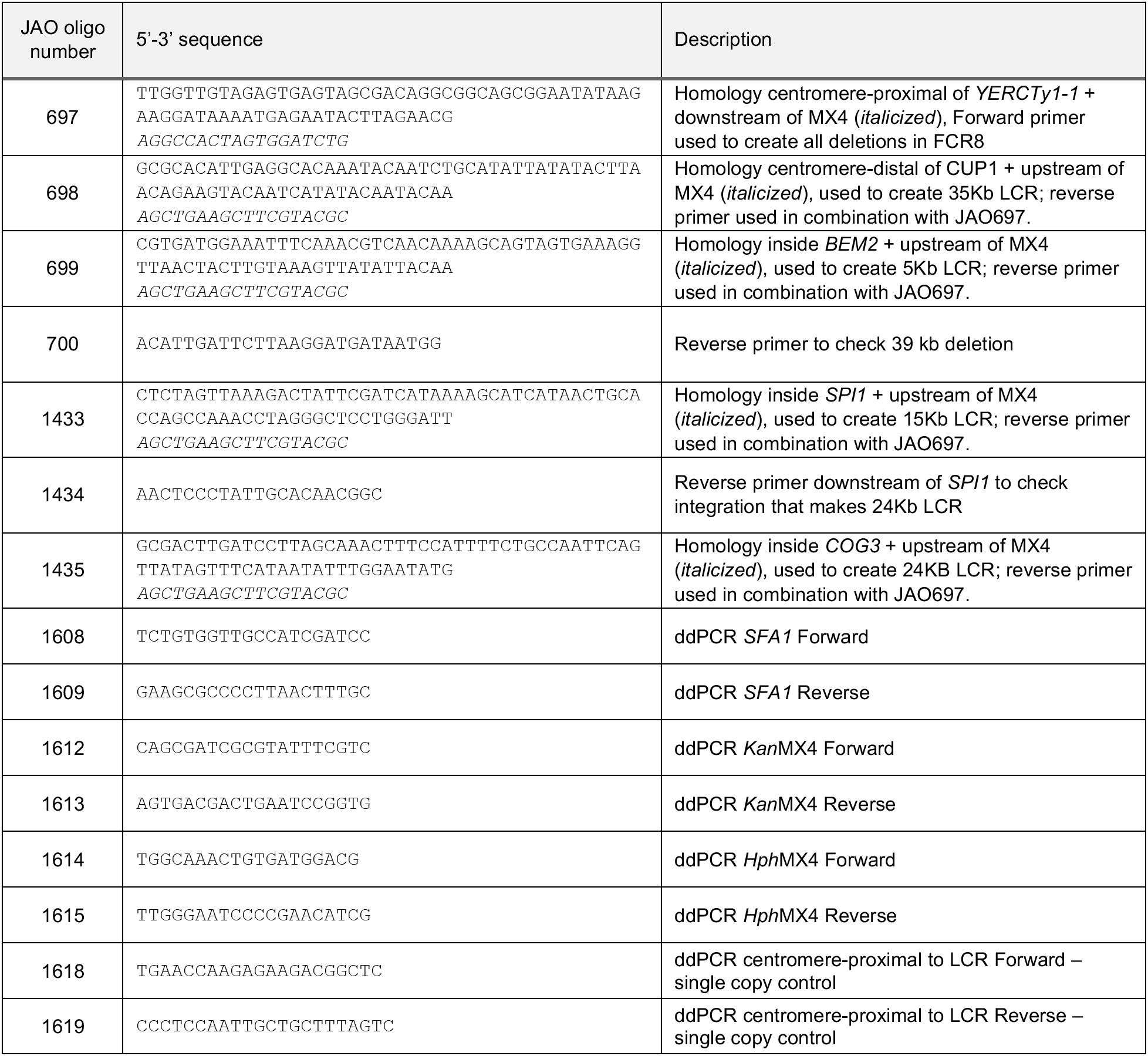
Oligonucleotide primers used in this study.

## Notes

### Competing Interest Statement

The authors have declared no competing interest.

